# Identification of a BACH1 lung cancer signature: A novel tool for understanding BACH1 biology and identifying new inhibitors

**DOI:** 10.1101/2025.01.10.632407

**Authors:** Donika Klenja-Skudrinja, David Walker, Kevin X. Ali, Maureen Higgins, Angana AH Patel, Dorota Raj, Anna Creelman, Charlotte McDowall, Tomasz Wenta, Erik Larsson, Clotilde Wiel, Volkan I. Sayin, Laureano de la Vega

## Abstract

The transcription factor BACH1 is a transcriptional repressor with a central role in regulating oxidative stress and anti-inflammatory pathways, emerging as a promising therapeutic target for multiple conditions, including neoplastic malignancies, neurodegenerative disorders, ischemia-reperfusion injuries and sickle cell disease. In the field of cancer BACH1 has gained significant attention, with BACH1 overexpression correlating with poor prognosis and metastasis across various cancer types; however, despite this increasing relevance of BACH1, no universal pro-metastatic mechanism or transcriptional signature for BACH1 has been identified which is a major limitation for this growing field. To address this, we performed RNA-Seq coupled with ChIP-Seq in BACH1-proficient and BACH1-deficient lung cancer cells, identifying a set of common BACH1 directly regulated genes, which we thoroughly validated in a large panel of cancer cells. This novel lung cancer BACH1 transcriptional signature is highly sensitive and specific to BACH1 perturbations (both genetic and pharmacological) and does not respond to NRF2 modulation, underscoring its specificity. This signature not only represents a robust surrogate for BACH1 activity, but we also provide evidence of its potential value as a tool to i) identify novel BACH1 inhibitors, and ii) provide insights into BACH1’s pro-metastatic role.

## Background

The transcription factor BACH1 (broad complex, tramtrack and bric à brac and cap’n’collar homology 1) is a widely expressed transcriptional repressor involved in antioxidant and anti-inflammatory response pathways^1–5^. BACH1 depletion activates the expression of several cytoprotective genes which explains why BACH1 appears as an attractive therapeutic target against a variety of inflammatory and oxidative stress-related conditions including Huntington’s^6^ and Parkinson’s disease^7^, sickle cell disease^8^, ischemia/reperfusion injury^9,10^, non-alcoholic steatohepatitis^11^, insulin resistance^12^, coronary artery disease^5,13^ and tuberculosis infections^14^.

A well-characterised counterpart for BACH1 is the transcriptional activator NRF2 (Nuclear factor erythroid 2-related factor 2). Both BACH1 and NRF2 bind to similar genomic sequences termed antioxidant response elements (AREs), sharing several target genes while exerting opposite roles^2^, i.e. BACH1 repressing and NRF2 activating their transcription. As they share a subset of target genes, BACH1 inhibition could present a similar profile and be mistaken for NRF2 activation. However, even though they recognise very similar sequences, BACH1 has unique targets that are not regulated by NRF2 (and *vice versa*), and the reason behind this specificity is not clear. BACH1 can also act as a transcriptional activator^15–21^ but the molecular mechanisms supporting its activator role are not well understood.

Seemingly unrelated to its role in inhibiting cytoprotective pathways, BACH1 has also emerged as an important pro-metastatic factor. BACH1 is overexpressed in various tumour types correlating with poor prognosis and recurrence^16–18,20,22^, it promotes cancer cell invasion (*in vitro*) ^15,16,18,20,22–24^ and metastasis (*in vivo*)^16,18,20,22^. Accordingly, BACH1 depletion impairs tumour spread in preclinical models of lung, breast, and pancreatic cancer among others^17,18,20,22^. While it appears that the pro-metastatic role of BACH1 is tumour-type agnostic, the most robust data comes from studies in lung cancer^25^.

A sensible approach to better understand the role of BACH1 is to identify its target genes. However, there is currently a lack of clarity about the genes directly regulated by BACH1 in different tissues. The reported BACH1 target genes vary based on the model and approach used ^3,7,17,20,22,26^ with the notable exception of the stress-inducible enzyme Heme oxygenase 1 (HMOX1)^1^, which consistently emerges as one of the most induced gene in response to BACH1 depletion. HMOX1 catalyses the first step of the oxidative degradation of the heme group to carbon monoxide, iron, and biliverdin and has potent antioxidant and anti-inflammatory properties. Nevertheless, the BACH1 pro-metastatic role appears to be multifactorial, and different studies have suggested various target genes driving BACH1 pro-metastatic role. These include genes involved in migration and invasion (such as MMP1 and CXCR4 in breast cancer^16^), metabolism (such as HK2 and GAPDH in lung cancer^18^) and genes involved in the epithelial to mesenchymal transition (such as CDH1 and FOXA1 in pancreatic cancer^20^ and CDH2, SNAI2 and VIM in oesophageal cancer^19^). At this point, no general pro-metastatic mechanism or transcriptional signature has been identified for BACH1.

While BACH1 has become a relevant factor in many disease areas, the BACH1 field is still in its early stages. Developing the field further requires robust tools and controls to study BACH1 biology, such as a clear understanding of the pathways regulated by BACH1, a panel of specific target genes that could be used as surrogates for BACH1 activity, and specific and well-characterised inhibitors and antibodies for diverse applications (e.g. western blots, immunohistochemistry and immunofluorescence). In this manuscript, we report the identification of a novel BACH1 lung cancer signature that is also conserved in many other cell types. This signature is sensitive (responds to both genetic and pharmacological perturbation) and specific for BACH1 (does not respond to NRF2 modulation) and can be used to provide further insight into the pro-metastatic role of BACH1 in lung cancer cells and as a tool to identify new BACH1 inhibitors.

## Material and Methods

### Cell Culture

Cells were cultured in RPMI 1640 Medium (GIBCO) (HaCaT, H1944, H460, H2228, H358, H1975, H1792, A2780, AsPC1, DLD1, HK2), Dulbecco’s Modified Eagle Medium (DMEM) (H1299, A549, MDA-MB-231, MDA-MB-468, MDA-MB-436, VCaP, U87, Breast Fibroblasts, Breast CAFs) or DMEM F12 (1:1) (lung fibroblasts) at 37 °C and 5% CO2. All media was obtained from Thermo Fisher Scientific and supplemented with 10% FBS. All cells were either validated by STR profiling. or obtained from ATCC, and were routinely tested for mycoplasma. CRISPR-edited BACH1-KO cells were produced as previously described^27,28^. In short: The endogenous *BACH1* gene, was edited by transfecting cells with either pLentiCRISPR-v2 (a gift from Dr Feng Zhang, Addgene plasmid #52961) or pLentiCRISPRv2-blast (#98293, Addgene) containing single-guide (sg) RNAs directed against *BACH1* (CGATGTCACCATCTTTGTGG and GACTCTGAGACGGACACCGA, CCACTCAAGAATCGTAGGCC or TACTCAGCCTTAATGACCAG); control cells, referred to wild type (WT), are the pooled population of cells transfected with an empty pLentiCRISPRv2 vector after treatment with the appropriate antibiotic (puromycin or blasticidin).

Reconstituted cells were obtained by transducing BACH1-KO cells with either Lentiviral BACH1-GFP-Puro or control GFP particles obtained from Origene.

### Reagents

Antibodies against BACH1 (F-9) and LAMIN B2 (C-20) were obtained from Santa Cruz Biotechnology. Antibody against ALPHA-TUBULIN was obtained from Sigma-Aldrich. HRP-conjugated secondary antibodies were obtained from Life Technologies. Dimethyl sulfoxide (DMSO) was from Sigma-Aldrich. Hemin, CDDO, CDDO-Me and CDDO-TFEA were obtained from Cayman Chemicals. R,S-sulforaphane (SFN) was purchased from LKT Laboratories. TBE56 was synthesized as described^29^. Paeoniflorin was obtained from Selleck Chemicals. All siRNAs used were OnTargetplus SMARTPool siRNAs (mixture of 4 siRNAs provided as a single reagent) obtained from Horizon Discovery.

### Cell Transduction

Cells were incubated with the media-containing virus complemented with Polybrene (8 μg/ml) for 16 hours, followed by a medium exchange. Transduced cells were selected by further growth in the presence of 4 μg/ml puromycin; the surviving cells were tested by immunoblotting for BACH1 expression.

### siRNA Cell transfections

On the day prior to transfection, cells were plated to the required cell density (70-90% confluency). Lipofectamine RNAiMAX (Invitrogen) was used. The siRNA and lipofectamine were individually diluted in Optimem (Gibco) and incubated for 10 minutes at room temperature. Diluted siRNA was added to the diluted Lipofectamine solution (1:1 ratio) and further incubated for 15 minutes. siRNA-lipid complex was added to the cells and incubated overnight in a humidified incubator at 37°C and 5% CO2. The next morning, the medium was replaced with fresh medium and cells were incubated 36 hours more before collecting them.

### Quantitative real time PCR (rt-qPCR)

RNA was extracted using GeneJET RNA Purification Kit (Thermo Fisher Scientific) and 500 ng of RNA per sample was reverse-transcribed to cDNA using Omniscript RT kit (Qiagen) supplemented with RNase inhibitor according to the manufacturer’s instructions. The resulting cDNA was processed using TaqMan Universal Master Mix II (Life Technologies, Carlsbad, CA, USA) as well as corresponding Taqman probes (Supplementary Material and Methods). Gene expression was determined using a QuantStudio 7 Flex qPCR machine by the comparative ΔΔCT method. All experiments were performed between two and seven times and data were normalized to the housekeeping gene HPRT1.

### Cell lysis and western blot

Cells were washed and harvested in ice-cold phosphate-buffered saline (PBS). For <u>whole cell</u> <u>extracts,</u> cells were lysed in RIPA buffer supplemented with phosphate and protease inhibitors. Lysates were sonicated for 20 s at 20% amplitude and then cleared by centrifugation for 10 min at 4 °C. For <u>subcellular fractionation,</u> cells were resuspended in 400 μl of low-salt buffer A (10 mM Hepes/KOH pH7.9, 10 mM KCL, 0.1 mM EDTA, 0.1 mM EGTA, 1 mM β-mercaptoethanol) and after incubation for 10 min on ice, 10 μl of 10% NP-40 was added and cells were lysed by gently vortexing. The homogenate was centrifuged for 30 s at 13,200 rpm, the supernatant representing the cytoplasmic fraction was collected and the pellet containing the cell nuclei was washed 4 additional times in buffer A. The pellet containing the nuclear fraction was then resuspended in 100 μl high-salt buffer B (20 mM Hepes/KOH pH7.9, 400 mM NaCl, 1 mM EDTA, 1 mM EGTA, 1 mM β-mercaptoethanol). The lysates were sonicated and centrifuged at 4 °C for 10 min at 13,200 rpm. The supernatant representing the nuclear fraction was collected. Protein concentration was determined using the BCA assay (Thermo Fisher Scientific, Waltham, MA, USA). Lysates were mixed with SDS sample buffer and boiled for 5 min at 95 °C. Equal amounts of protein were separated by SDS-PAGE, followed by semidry blotting to a polyvinylidene difluoride membrane (Thermo Fisher Scientific). After blocking the membrane with 5% (w/v) non-fat dried milk dissolved in Tris buffered saline (TBS) with 0.1% v/v Tween-20 (TBST), membranes were incubated with the primary antibodies overnight at 4°C. Appropriate secondary antibodies coupled to horseradish peroxidase were detected by enhanced chemiluminescence using ClarityTM Western ECL Blotting Substrate (Bio-Rad, Hercules, CA, USA). The resulting protein bands were quantified and normalised to each lane’s loading control using the ImageStudio Lite software (LI-COR). For whole cell extracts, the protein of interest was normalised against LAMIN or TUBULIN. LAMIN was used as an internal control for nuclear extracts and TUBULIN was used as controls for cytoplasmic extracts.

### Cell viability assay

Alamar Blue (Thermo Fisher Scientific) was used to determine cell viability after drug treatment. Cells were seeded in 96-well plates to 50–60% confluency and treated the next day with the corresponding compounds for 48 hours. After treatment, Alamar Blue was added to the wells (1:10 ratio) and after six hours of incubation at 37 °C the fluorescence was measured (excitation 535 and an emission at 590 nm) using a microplate reader (Infinite F Plex Tecan). Viability was calculated relative to the DMSO-treated control.

### Transwell Migration Assay

The transwell migration assay was conducted using Corning^TM^ Transwell^TM^ Multiple Well Plate with 6.5 mm inserts and 8.0-μm pore permeable polyester membrane (Fisher Scientific). Cells were trypsinised and were then washed with PBS before resuspending them in serum-free media. For migration assay of the siRNA transfected A549 cells, 150 ul containing 70000 cells in serum-free media were seeded on the upper chamber of the insert and 600 ul of complete media (10% FBS v/v) was added to the bottom chamber.

After 16h, the transwells were washed in PBS before using 4% PFA to fix the migrated cells for 15 minutes, followed by another PBS wash and finally staining with crystal violet (0.025%) for 20 minutes. The crystal violet was then aspirated and the cells that did not migrate and remained in the upper chamber were removed using a cotton swab wet in PBS. The transwells were washed to remove excess crystal violet by dipping them a few times in a beaker filled with distilled water. Imaging of the transwells was conducted on five different areas of the well under bright-field microscope (20x). Images were analysed blind to the experimental conditions, and stained cells were automatically counted using ImageJ. The mean cell count was derived from five images per well and normalized to respective controls.

### Chromatin Immunoprecipitation-Sequencing (ChIP-Seq)

Chromatin immunoprecipitation (ChIP) experiments were performed on A549 and A549 *BACH1* knockout cells using the BACH1 antibody (14018-1-AP, ProteinTech). Library preparation and NGS sequencing were performed by Active Motif. Sequencing was performed on an Illumina NextSeq 500 platform, generating 75 bp single-end reads. The reads were aligned to the human genome (hg38) using the BWA algorithm, retaining only high-quality reads with ≤2 mismatches and unique genomic alignments. Fragment densities were calculated by extending aligned reads in silico to 200 bp and binning the genome into 32-nt intervals. Peaks corresponding to BACH1 binding sites were identified using MACS2 and normalized for tag counts across samples. Genomic intervals were annotated for nearby genes and visualized as bigWig files in the Integrative Genomics Viewer (IGV, v.2.18.4)^30^.

Standard normalization was performed for comparative analysis by down-sampling usable tags across samples in each group to match the sample with the fewest tags. Peaks were called using either the MACS2 (p-value cutoff of 1e^-7^ for narrow peaks and 1e^-1^ for broad peaks) or SICER (FDR 1e^-10^ with a 600 bp gap parameter) algorithms. False peaks were filtered using the ENCODE blacklist. Known motifs and *de novo* motifs were identified using HOMER’s *findMotifsGenome* tool with default parameters and sequences within ±200 bp of the center of the top 2500 peaks^31^. Find Individual Motif Occurrences (FIMO) with default parameters (MEME v.5.5.7) was used to scan the sequences for individual matches to the BACH1 and NRF2 motifs^32^.

To identify and compare NRF2 binding sites with BACH1 binding sites in A549 cells, publicly available ChIP-Seq data was obtained from NCBI Gene Expression Omnibus (accession number: GSE141497)^33^. The reads were aligned to the human genome (hg38) and processed as bigWig files to identify the overlapping binding regions visualized in IGV.

### RNA Sequencing (RNA-Seq)

A549 WT and *BACH1* KO cells were seeded in triplicate and 2 million cells from each condition were processed for the RNA sequencing. Library preparation and sequencing were done at Active Motif. Sequencing reads generated by Illumina sequencing were aligned to the ENSEMBL GRCh38 reference genome using STAR RNA-Seq aligner^34^ using default parameters. Genes with zero counts were excluded. Pairwise comparisons were performed with the DESeq2 software package^35^. A Wald test was used to determine significance for comparisons with replicates, and any gene with an adjusted p-value of less than 0.1 was considered differently expressed. Differentially expressed genes with a shrunken log fold change greater than 0.3 were considered for comparisons without replicates. Publicly available microarray data from the DMSO—or paeoniflorin (PF)—treated human pancreatic cancer cell line Capan-1 from NCBI Gene Expression Omnibus (accession number GSE97124)^36^ was used to identify significantly altered genes in response to PF treatment.

### Integration of ChIP-Seq and RNA-Seq Data

To identify genes with both high BACH1 binding affinity and significant differential expression upon *BACH1* knockout, ChIP-Seq and RNA-Seq datasets were integrated. For the ChIP-Seq data, all peaks within genes were included, along with any peak located within 10 kb upstream or downstream of genes. To priorities high-affinity binding sites, genes with peak values (Log₂ Ratio BACH1/BACH1 KO) below 0.7 were excluded. The RNA-Seq dataset included differentially expressed genes, characterized by two key metrics: Log2 Fold Change (reflecting the magnitude of expression differences between A549 WT and BACH1 KO cells) and adjusted p-value (padj). To integreate the datasets, a combined score was calculated for each gene, with weights assigned to the metrics as follow: Combined Score = (0.5 × Log₂ Ratio) + (0.25 × |Log₂ Fold Change|) + (0.25 × −log₁₀(adjusted p-value)) This formula ensured a balanced contribution from both ChIP-Seq and RNA-Seq data, with 50% weight assigned to the ChIP-Seq Log2Ratio and 50% to RNA-Seq metrics (25% for the magnitude of fold change and 25% for statistical significance). Gene Ontology (GO) enrichment analysis was performed to identify enriched Biological Processes (BP) and Molecular Functions (MF) associated with the input gene list derived from the integrated ChIP-Seq and RNA-Seq data. The analysis was conducted using R version 4.4.1 within the RStudio environment. Bioconductor packages clusterProfiler^37^, org.Hs.eg.db, enrichplot, dplyr, and ggplot2 were employed for gene annotation, enrichment analysis, and high-quality visualizations.

### Statistical analysis

Experiments were repeated 2-7 times. Data were analysed using Graphpad Prism statistical package using t-tests to test for differences between control and experimental treatments. All results are presented as mean ± SD unless otherwise mentioned. *P ≤ 0.05, **P ≤ 0.01, ***P ≤ 0.001.

## Results

### Identification of common BACH1 target genes in lung cancer cells

We compiled a list of 30 published BACH1 target genes identified in cancer cells (list of genes provided in Supplementary Material and Methods) and compared their expression between WT and BACH1-KO cells in two lung (A549 and H1299) and two breast (MDA-MB-231 and MDA-MB-468) cancer cell lines. Our analysis reveals minimal overlap between BACH1-regulated genes across the four cell lines (Fig. 1A and Suppl. Fig. S1A) with only HMOX1 being consistently regulated more than two-fold in all cell lines. This highlights our lack of knowledge of general, cell type independent, BACH1 targets. The relevance of BACH1 in lung cancer has been clearly demonstrated; however, to deepen our understanding of BACH1’s biology and its links to lung cancer metastasis, a clear understanding of the genes and pathways regulated by BACH1 in human lung cancer cells is still necessary. To identify a robust panel of BACH1 target genes in lung cancer cells we performed integrative RNA-Seq and ChIP-Seq analyses in BACH1 proficient (WT) and deficient (BACH1-KO) lung adenocarcinoma A549 cancer cells (Fig. 1B-E, Suppl. Fig. S1 and Suppl. Files). ChIP-Seq analysis identified significant BACH1 binding peaks across the genome (Fig S1B), with enrichment around transcription start sites (TSS), consistent with BACH1’s role as a transcription factor (Fig. 1B). *De novo* motif analysis showed that 74.39% (p-value = 1e-12798) of the identified targets were associated with TGACTCAGC consensus sequence for BACH family transcription factors shared by BACH1 and BACH2 (Fig. 1C). Furthermore, the top known motif identified in 57.82% of the targets (p-value = 1.0e-14620) is the canonical BACH1 motif, further validating our ChIP-Seq results (Fig. 1C and Suppl. Fig. S1C and Suppl. Files). By integrating the ChIP-Seq and RNA-Seq results, we obtained a list of genes that were both directly bound and significantly regulated by BACH1 (Fig. 1D, 1E and Supp Files). Among these, *HMOX1, HTRA3, ZNF469, NRCAM and FTH1* emerged as the top five genes that were directly bound by BACH1 and significantly up-regulated in BACH1-KO cells. Gene Ontology (GO) functional annotation of these genes revealed significant enrichment in biological processes (Fig. 1F) and molecular functions (Suppl. Fig. S1D) related to cell adhesion, cell migration and cytoskeleton remodelling, consistent with BACH1’s role in cancer metastasis. To further investigate BACH1’s regulatory functions, we next analysed its binding sites identified in A549 cells (Fig. 1G *in blue*). Additionally, using publicly available NRF2 ChIP-Seq data from A549 (GSE141497), we observed that NRF2 binds each BACH1-bound region (Fig. 1G *in red*). This analysis also revealed distinct binding profiles among the five target genes, characterised by variations in the number and arrangement of putative binding motifs. For example, a single BACH1 binding region containing one shared motif for BACH1 and NRF2 was identified for *FTH1* gene. In contrast, *HMOX1* and *NRCAM* displayed multiple binding regions, each containing one or more BACH1 motifs. Meanwhile, *ZNF469* and *HTRA3* displayed multiple BACH1 binding motifs clustered within a single region. Notably, NRF2 motifs did not consistently overlap with BACH1 motifs, suggesting that these transcription factors may employ diverse binding/competing strategies for the different loci.

**Figure 1.**
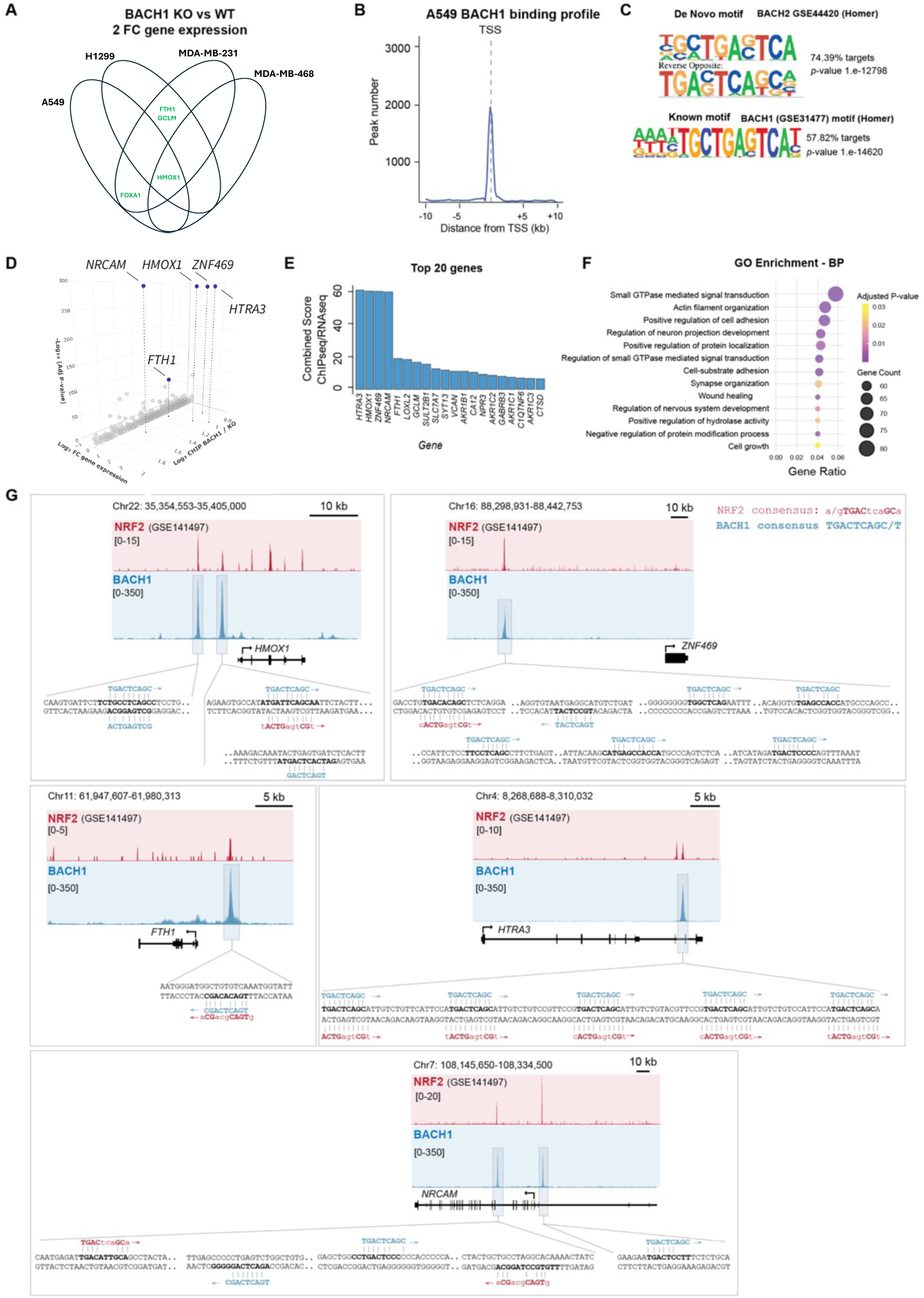
Comprehensive analysis of BACH1 binding and functional impact. **A)** Venn diagram showing the overlap of differentially expressed BACH1 target genes (2-fold cutoff) upon BACH1 depletion across two lung cancer cell lines (A549 and H1299) and two breast cancer cell lines (MDA-MB-231 and MDA-MB-468). **B)** BACH1 binding profile showing the distribution of peak numbers relative to the transcription start site (TSS) from the ChIP-Seq analysis. **C)** Sequence logos of BACH motifs. The de novo motif (BACH2 GSE44420) matches 74.39% of targets (p-value = 1.e-12798). The known motif, BACH1 motif (GSE31477), matches 57.82% of targets (p-value = 1.e-14620). **D)** 3D Scatter plot showing the relationship between gene expression (RNA-Seq Log₂ fold change), adjusted *p*-value (RNA-Seq -Log₁₀) and ChIP-Seq Binding Peak value (Log₂ Ratio BACH1 WT / BACH1 KO). A cutoff of 0.7 was applied to the ChIP-Seq binding ratio to exclude low-affinity targets, ensuring that only genes with significant binding (either within 10 kb of TSS or in gene) were considered. Highlighted genes are the top 5 genes *NRCAM1, HMOX1, ZNF469, HTRA3,* and *FTH1*. **E)** Bar graph showing the top 20 genes ranked by their combined ChIP-seq and RNA-seq scores. **F)** Dot plot showing enriched Gene Ontology (GO) Biological Processes (BP) performed on data integrating RNA-Seq and ChIP-Seq genes. **G)** ChIP-Seq peak locations for BACH1 and NRF2 binding at the loci of the top 5 target genes (*HMOX1, ZNF469, FTH1, HTRA3,* and *NRCAM*). Tracks showing BACH1 and NRF2 binding profiles are displayed in blue and red, respectively, highlighting overlapping and distinct binding regions. Note that additional NRF2 binding motifs in the regions of interest are not displayed for *HMOX1* and *ZNF469*.

### Validation of the BACH1 signature

We next conducted a thorough validation of the 5-gene BACH1 signature in A549 cells to ensure that the genes were bona fide BACH1 targets. This validation included comparing WT and BACH1-KO cells generated with 3 different gRNAs in two different A549 parental cells (from two different laboratories) (Fig. 2A) and using siRNAs against BACH1 as an orthogonal method (Fig. 2B). These experiments validated the identified genes as *bona fide* BACH1 target genes. Additionally, we also performed reconstitution experiments introducing back BACH1-GFP into A549 BACH1-KO cells. Although the levels of nuclear BACH1 in the reconstituted cells were lower than in the WT cells, the partially reconstituted BACH1 cells still showed an opposite pattern of gene regulation for the identified genes (i.e., lower levels in the BACH1 reconstituted cells) supporting the role of BACH1 as a repressor for these genes (Fig. 2C). As previously showed (Fig. 1A), the lack of BACH1 target genes conservation across different cancer types and even between cell lines derived from the same cancer type is a significant limitation. To test the robustness of our signature, we measured the induction of the signature genes in response to BACH1 depletion in a panel of six lung cancer cell lines (Fig. 2D and validation of the cell lines in Suppl. Fig. S2A). These cell lines, which harbour different oncogenic drivers (e.g. A549, mutant KRAS; H1299, mutant NRAS; H1944, mutant KRAS; H2228, WT KRAS; H460, mutant KRAS; H358 mutant KRAS; and H1795, WT KRAS) and distinct NRF2/KEAP1 statuses (A549, mutant KEAP1; H1299, WT KEAP1; H1944, mutant KEAP1; H2228, mutant KEAP1; H460, mutant KEAP1; H358, WT KEAP1; H1795, WT KEAP1) demonstrated that the signature performs well in a variety of lung cancer cells. Of the identified genes, *HMOX*1, *ZNF469* and *HTRA3* were highly upregulated in most cell lines (H358 and H1795 did not express HTRA3 to detectable levels), whereas NRCAM and FTH1 were upregulated at varying levels. We also compared the expression of these genes between WT and BACH1-KO in the non-cancerous keratinocyte cell line HaCaT (Suppl. Fig. S2B), showing a pattern consistent with that previously identified in lung cancer cells. As *FTH1* is a well-known NRF2 target gene and its combined score (Fig. 1E) was clearly lower than that of the other top genes, we excluded it from subsequent experiments to enhance the specificity of the signature without compromising sensitivity. To further confirm the validity of the signature we also reconstituted two other BACH1-KO cell lines (HaCaT and H1944) with BACH1-GFP and tested the expression of the 4 top target genes (Suppl. Fig. S2C and S2D), confirming that all genes except for *NRCAM*, were negatively regulated by BACH1. Although *NRCAM* expression is induced by BACH1 depletion in most cell lines tested (Suppl. Fig. S2E), its expression was recovered in only one of three BACH1 reconstituted cell lines (Fig. 2C, Suppl. Fig. S2C and S2D). Despite strong indications that it is a genuine BACH1 target, our stringent experimental approach could not provide conclusive confirmation, resulting in its exclusion from subsequent experiments.

**Figure 2.**
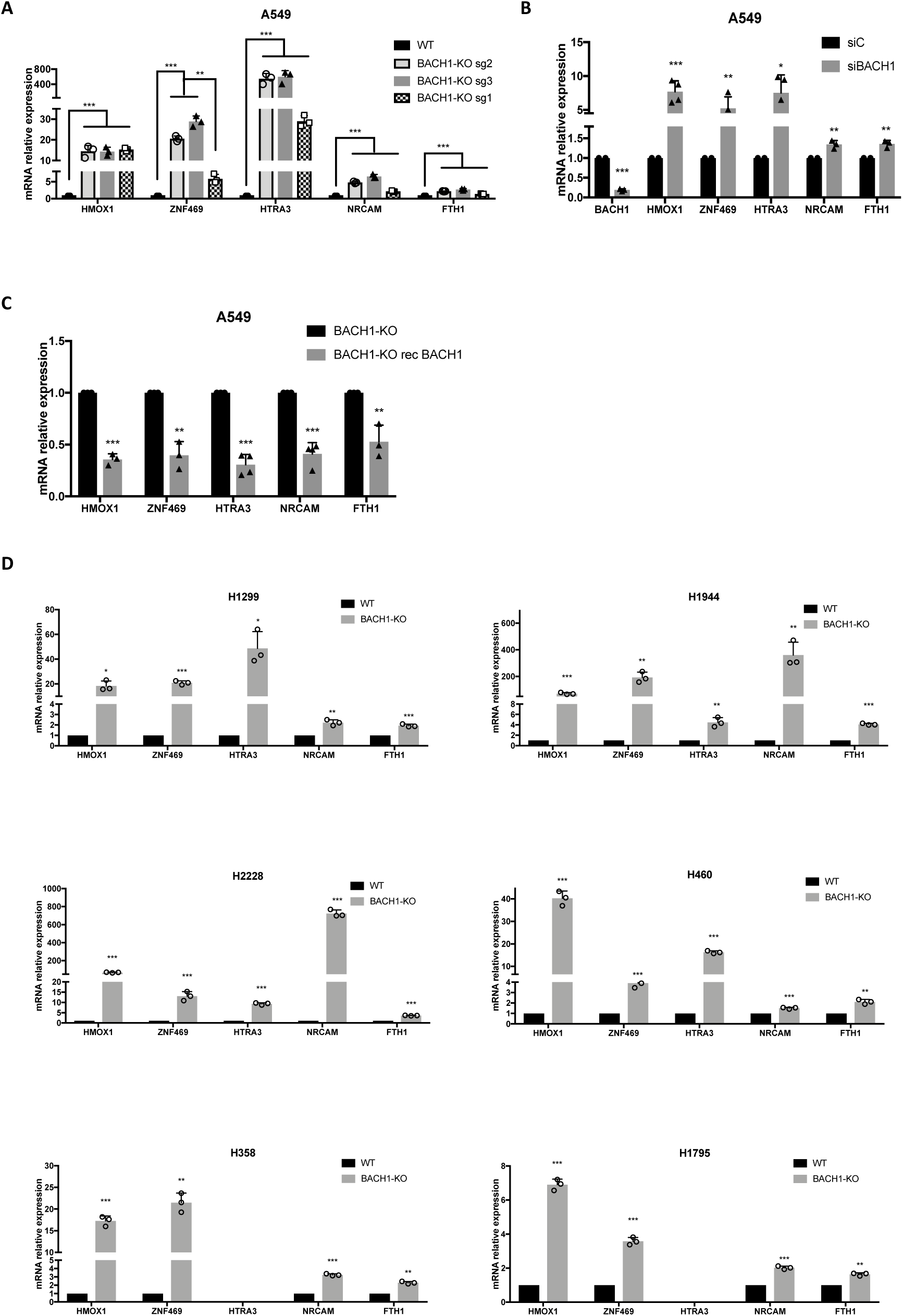
Validation of the BACH1 signature in lung cancer cells. **A)** A549 control cells (WT) and BACH1-KO cells produced with three different gRNAs were harvested and lysed, and mRNA levels of the indicated genes were analysed by real-time qPCR. Data (n=3) represent means± SD and are expressed relative to the WT sample. **B)** A549 cells were transfected with either siControl (siC) or siBACH1. 48 hours later cells were lysed, and mRNA levels of the indicated genes were analysed by real-time qPCR. Data (n= 3-5) represent means± SD and are expressed relative to the siControl sample. **C)** A549 BACH1-KO cells were infected with either GFP control or with BACH1-GFP virus (rec BACH1) and selected with puromycin (4 µgr/ml). 72 hours later cells were lysed, and mRNA levels of the indicated genes were analysed by real-time qPCR. Data (n=3-5) represent means± SD and are expressed relative to the BACH1-KO (GFP control) sample. **D)** A panel of six control (WT) and BACH1-KO lung cancer cell lines were compared for their expression of the indicated genes. Data (n=3) represent means± SD and are expressed relative to the control sample.

Using the reduced three-gene signature, we next measured their induction in response to BACH1 depletion using siRNAs in a large panel of cancer and non-cancerous cells (Suppl. Fig. S2F). The signature works in most cells (but not in all), demonstrating that induction of *HMOX1, ZNF469* and *HTRA3* is a strong surrogate for BACH1 depletion in lung cancer cells and a good pan-tissue BACH1 signature.

As BACH1 and NRF2 are known to have many common target genes and ChIP-Seq data indicate that NRF2 can also bind to our three signature genes (Fig 1G), we tested whether the newly identified signature was specific for BACH1 depletion or would also reflect an NRF2 stabilisation signature. We compared the expression of *HMOX1*, *HTRA3* and *ZNF469* in response to either BACH1 depletion, KEAP1 depletion or dual BACH1 and KEAP1 depletion. KEAP1 is the main E3-ligase controlling NRF2 stability, and KEAP1 depletion/inhibition results in a strong NRF2 stabilisation and a subsequent induction of NRF2 target genes. To control for KEAP1 depletion and NRF2 stabilisation, we included in the analysis the bona fide NRF2 target gene *AKR1B10*^6,38^. We chose the lung cancer cell line H1299 for these experiments due to its high BACH1 levels and functional KEAP1 (leading to low NRF2 levels), making it an ideal model for studying both BACH1 and KEAP1 depletion. Our data show that *HMOX1, ZNF469* and *HTRA3* are significantly induced by siBACH1 but barely change in response to siKEAP1, showing that, at least in this cell line, the expression of these three genes is not significantly regulated by NRF2 induction, and therefore confirming their induction as a specific proxy for BACH1 depletion (Fig. 3A). Given the known role of NRF2 regulating HMOX1, this might appear contradictory; however, we have previously shown that in many human cell lines, particularly in the context of high BACH1 levels, NRF2 activation often results in a very limited *HMOX1* induction^6,39,40^.

**Figure 3.**
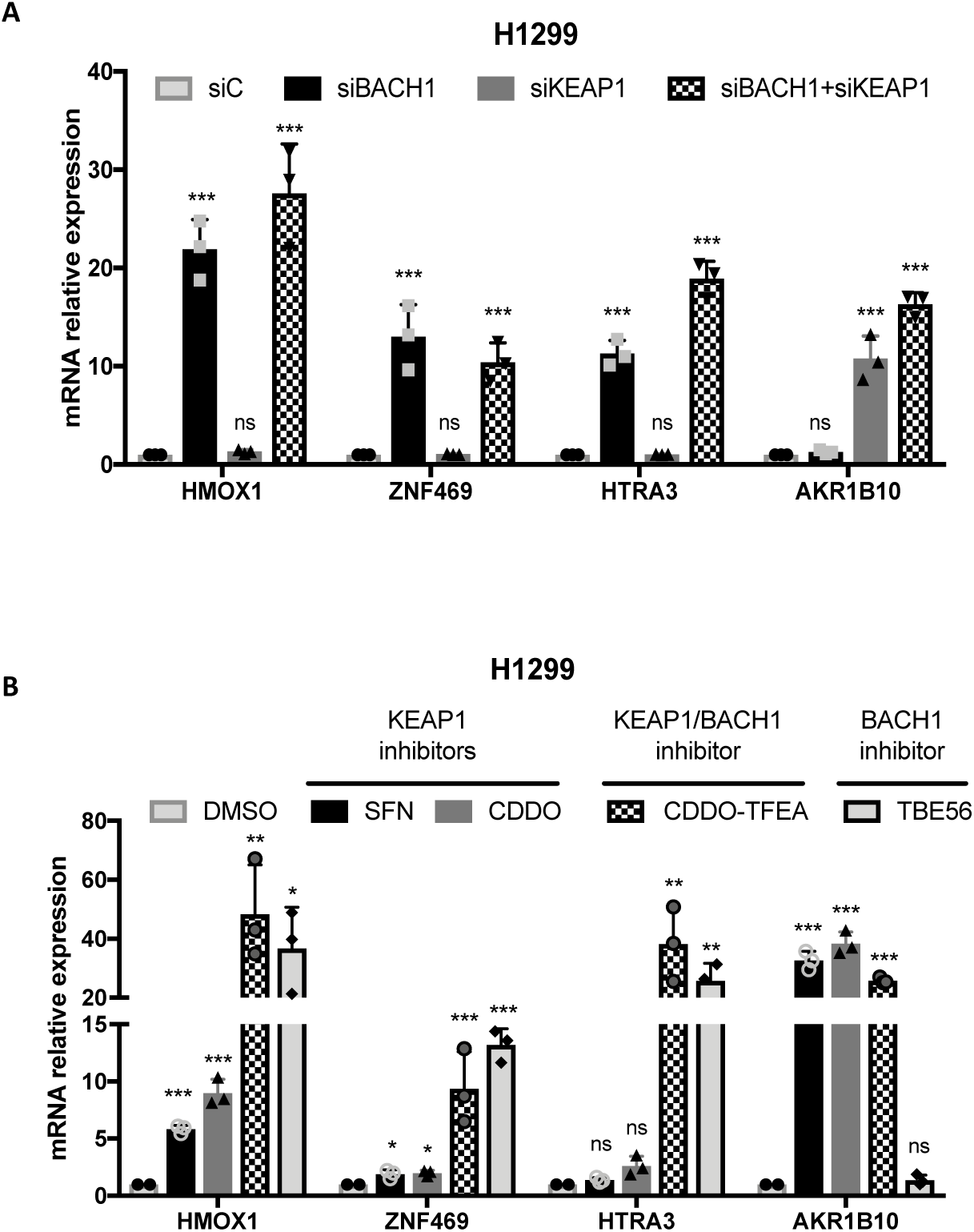
Induction of HMOX1, ZNF469 and HTRA3 is a specific surrogate for BACH1 depletion/inhibition. **A)** H1299 cells were transfected with either siControl (siC), siBACH1, siKEAP1 or both siBACH1 and siKEAP1. 48 hours later cells were lysed, and mRNA levels of the indicated genes were analysed by real-time qPCR. Data (n=3) represent means± SD and are expressed relative to the siControl sample. **B)** H1299 cells were treated with either DMSO (0.1%, v/v), SFN (5 μM), CDDO (100 nM), CDDO-TFEA (100 nM) or TBE56 (100 nM) for 16 hours. After that, cells were lysed, and mRNA levels of the indicated genes were analysed by real-time qPCR. Data (n=3) represent means± SD and are expressed relative to the DMSO treated sample.

Additionally, we also tested whether the signature was sensitive to BACH1 pharmacological inhibition and whether, as suggested by the genetic approach, the signature could discriminate between BACH1 inhibitors and KEAP1 inhibitors (NRF2 activators). To test this, we used the well-validated KEAP1 inhibitors SFN and CDDO, the BACH1 inhibitor TBE56^28^ and the dual KEAP1/BACH1 inhibitor CDDO-TFEA^27^ in the lung cancer cell line H1299. Figure 3B shows that the BACH1 inhibitors effectively induced the three genes of the signature, while KEAP1 inhibitors failed to do so, confirming that induction of these three genes is a specific and sensitive proxy for both BACH1 depletion and inhibition. Similar results were obtained in other cell lines (Suppl. Fig. S3A and S3B).

### Use of the signature to identify novel BACH1 inhibitors

We hypothesised that by interrogating transcriptomics databases against our signature we could identify compounds that inhibit BACH1. To test this hypothesis, we searched the GeoDatasets looking at compounds that induce the expression of HTRA3, ZNF469 and HMOX1. Based on this, the first study we identified was a transcriptomics study done in pancreatic cancer cells with the compound paeoniflorin^36^. Paeoniflorin is a natural compound derived from the root of the *Paeonia lactiflora* plant and is commonly used in herbal and traditional Chinese medicines. Paeoniflorin has a wide range of pharmacological properties, including analgesic antioxidant, anti-inflammatory and neuroprotective, with various described targets, including KEAP1/NRF2^36,41^. In the mentioned study, the most upregulated genes in response to paeoniflorin treatment were *NRCAM*, *HMOX1*, *HTRA3* and *ZNF469*, with *FTH1* also significantly upregulated. Based on the similarity between the transcriptomic profile upon paeoniflorin treatment and the one obtained after knocking out *BACH1* (Fig 4A), we hypothesised that paeoniflorin could be a BACH1 inhibitor. After an initial cell viability analysis to identify the concentrations of paeoniflorin that did not induce cell death (Suppl. Fig. S4A), we tested whether this compound reduced BACH1 levels in lung and breast cancer cell lines. Our results showed that paeoniflorin, at similar doses as those used in the transcriptomic analysis (which do not reduce cell viability), reduced the levels of BACH1 (total, nuclear and cytoplasmic levels) in lung and breast cancer cell lines (Fig 4B and Suppl. Fig. S4B and S4C). A time-dependent analysis showed that paeoniflorin reduced BACH1 protein levels within the 6-24 h time frame (Fig 4D).

**Figure 4.**
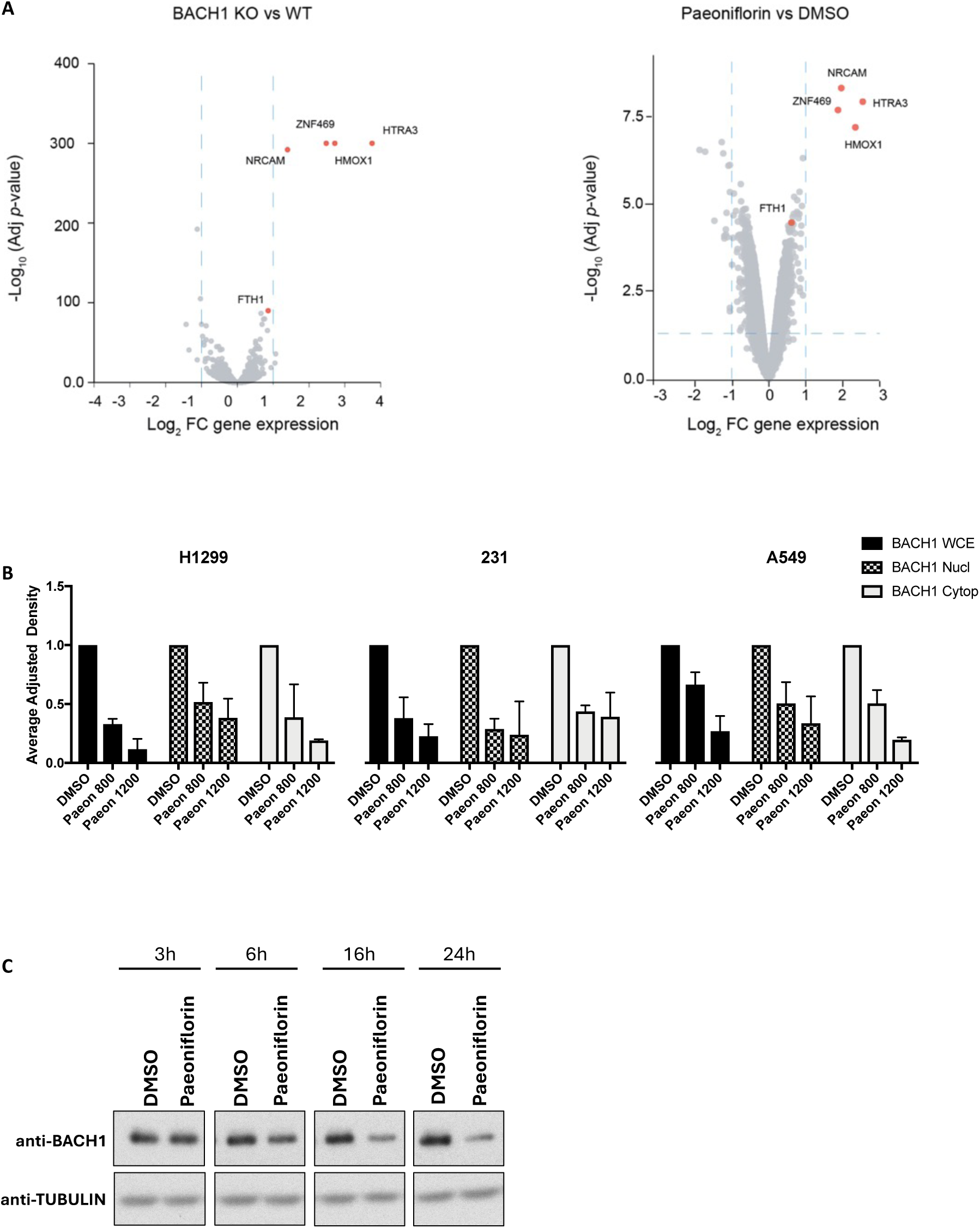
Validation of paeoniflorin as a novel BACH1 inhibitor. **A)** Volcano plot of RNA-Seq analysis comparing gene expression in A549 BACH1 wild-type (WT) vs. knockout (KO) cells (left panel) and DMSO vs. paeoniflorin treated Capan1 cells (right panel). Log₂ fold change (FC) in gene expression is plotted against the -Log₁₀ adjusted p-value. Highlighted genes (red) include ZNF469, NRCAM, HMOX1, HTRA3, and FTH1. **B)** Lung cancer cells (A549 and H1299) or breast cancer cells (MDA-MB-231) were treated with vehicle (DMSO) or with paeoniflorin at the indicated concentrations. After 16 hours cells were lysed either for whole cell extract (WCE) or were subjected to subcellular fractionation and separated in nuclear and cytoplasmic fractions. The figure shows the quantification of BACH1 protein levels normalized for TUBULIN or LAMIN levels; data (n=3) represent means± SD and are expressed relative to the DMSO-treated samples. Representative western blots shown in Suppl. fig. S4B and S4C **D)** H1299 cells were treated with vehicle (DMSO) or with paeoniflorin (1000 μM) for 3, 6, 16 and 24 hours. After that, cells were lysed and samples were analysed by western blot with the indicated antibodies.

### Relevance of the identified BACH1 target genes in its pro-metastatic role

In addition to its relevance as a tool to identify/validate BACH1 inhibitors, we used the information obtained from integrating our RNA-Seq and ChIP-Seq to obtain further insight into the biology of BACH1 in lung cancer. From the validated three gene signature we focused on the two newly identified BACH1 target genes, *HTRA3* and Z*NF469,* both of which are highly induced by BACH1 depletion in lung cancer cells (Fig. 5A and S5A). *HTRA3* (High temperature requirement A, Serine Peptidase 3) is a serine protease involved in matrix remodelling^42,43^ while *ZNF469* (Zinc-finger protein 469) regulates the expression of extracellular matrix components and its mutations have been linked to brittle cornea syndrome^44,45^. A previous report suggested that A549 cells did not express HTRA3^42^, but we confirmed that although the HTRA3 protein levels were very low in A549 WT cells, they were readily detected in A549 BACH1-KO cells (Fig. 5B)

**Figure 5.**
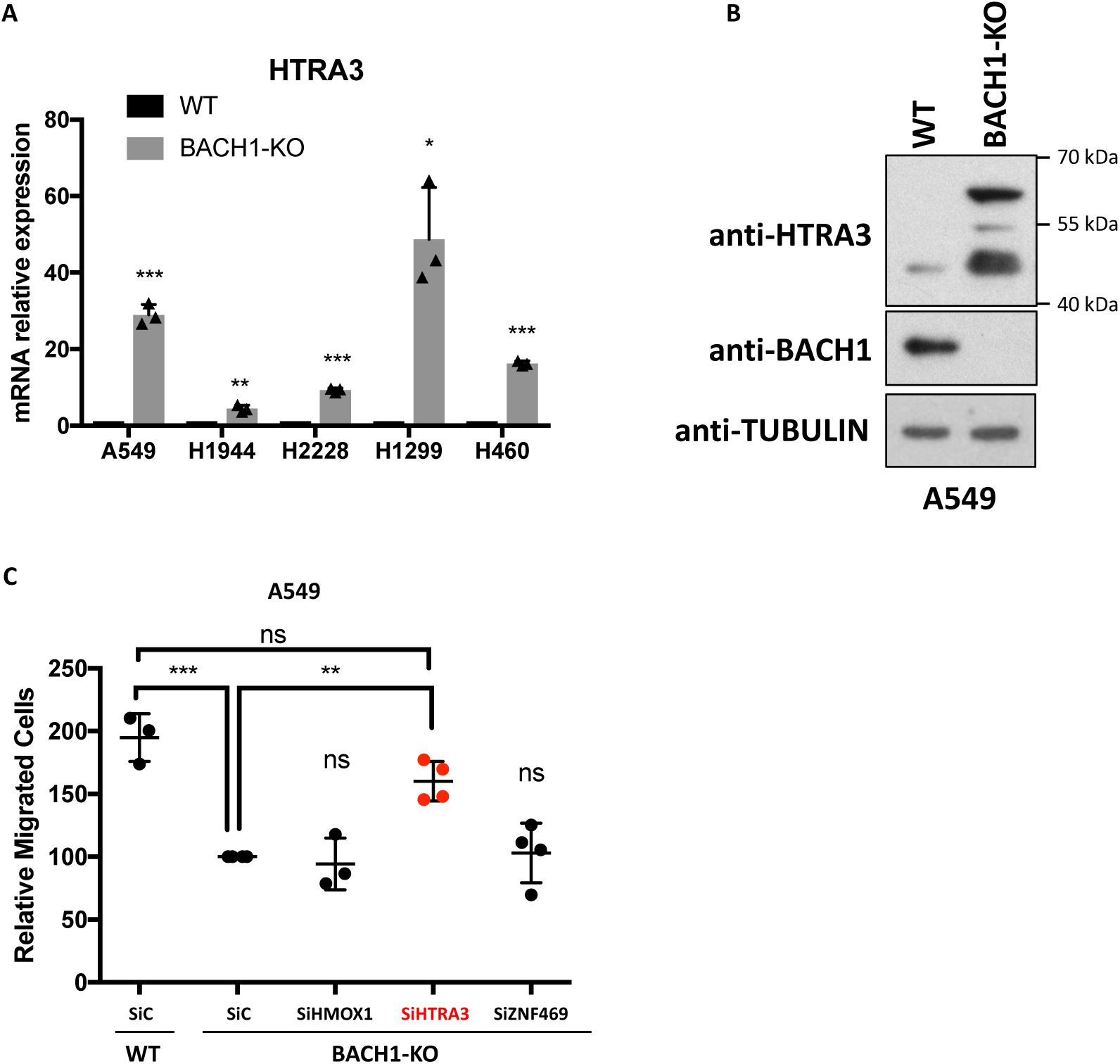
HTRA3 induction is necessary for the reduced cell migration in BACH1-KO lung cancer cells. **A)** A panel of control (WT) and BACH1-KO lung cancer cell lines were compared for their expression of *HTRA3*. Data (n=3) represent means± SD and are expressed relative to the control sample. **B)** Whole cell lysates from A549 control (WT) and BACH1-KO cells were compared for the levels of the indicated proteins, via western blot. **C)** A549 control cells transfected with siControl (siC) and A549 BACH1-KO cells transfected with either siControl (siC), siHMOX1, siHTRA3 or siZNF469 were compared for their migration capacity using transwell assays. Data (n=3-4) represent means± SD and are expressed relative to the BACH1-KO siControl sample. Transfection efficiency is shown in Suppl. Fig. S5B.

As BACH1 depletion/inhibition leads to reduced lung cancer cell migration^18,27,28^ we investigated whether induction of the signature genes was causally linked to the reduced migration in BACH1-KO cells. To test this, we silenced *HTRA3, ZNF469* or *HMOX1* in BACH1-KO cells and studied their migration. Neither the silencing of *HMOX1* nor *ZNF469* affected cell migration in BACH1-KO A549 cells; however, silencing *HTRA3* in BACH1-KO lung cancer cells (which have high levels of HTRA3 and reduced migration) largely recovered their migration (Fig 5C and Suppl. Fig. S5B), suggesting that HTRA3 upregulation is necessary for the reduced migration of BACH1-KO cells.

## Discussion

In this manuscript, we discuss the identification of a novel, specific and sensitive BACH1 lung cancer signature, which is also relevant in other tissue types.

Our analysis of 30 published BACH1 target genes shows little overlap of regulated genes in response to BACH1 depletion across four cancer cell lines. This suggests that BACH1 regulates distinct transcriptional programs in different cells, highlighting the importance of studying BACH1 biology in each specific context as its targets may differ significantly across tissue/cell types. To our knowledge, this is the first time a comprehensive integration of RNA-Seq and ChIP-Seq in human lung cancer cell lines using CRISPR-mediated knockouts has been performed to study BACH1. As expected, among the list of genes identified in the RNA-Seq to be negatively regulated by BACH1, were several known NRF2 target genes (e.g *GCLM, AKR1B1, SLC7A11, FTH1 and HIPK2*^46^). Furthermore, our analysis of the BACH1 and NRF2 binding sites in the identified genes showed overlapping peaks for NRF2 and BACH1 suggesting that both factors can bind to the same region, even if not the same motif. This interplay between BACH1 and NRF2 is also supported by the similarity between the NRF2 and BACH1 binding motifs, including the de novo BACH1 binding motif identified in our analysis.

Our identification of robust and specific surrogate genes for BACH1 inhibition is very valuable, as it can be combined with specific surrogate genes for KEAP1 inhibition to distinguish between BACH1, KEAP1, and dual KEAP1/BACH1 inhibitors. This differentiation is crucial as these inhibitors exhibit very similar profiles and could otherwise be mistakenly grouped. Identifying whether a compound specifically targets KEAP1, BACH1 or both is important as they will activate different transcriptional programs, and thus have different activity profiles and therapeutic indications.

Additionally, this signature can be used to validate novel BACH1 inhibitors or, in an unbiased manner, to identify unrecognised BACH1 inhibitors through transcriptomics. We showed that by mining transcriptomics databases for compounds with profiles matching our signature, we identified the natural compound Paeoniflorin as a novel BACH1 inhibitor, demonstrating the value of the signature as a useful discovery tool.

The identified BACH1-regulated genes (direct or indirectly regulated) provide further insight into the biology of BACH1 and its regulatory networks. As BACH1 has been involved in various processes including oxidative stress responses, heme and iron homeostasis, metabolism and cell migration/invasion, an understanding of the genes and pathways regulated by BACH1 in lung cancer cells is crucial to clarify its context dependent role. Using our signature, we identified HTRA3 as a potential novel effector of BACH1’s pro-migratory effect in human lung cancer cells. Various studies have highlighted a link between HTRA3 and lung cancer, with HTRA3 being downregulated in lung tumour tissue^47^, and higher HTRA3 levels correlating with lower recurrence and longer disease-free survival in lung cancer patients^47,48^. Furthermore, HTRA3 restricts lung cancer cell migration and invasion^43,47,48^ further supporting the new link we established between the reduced migration observed in BACH1-depleted cells and HTRA3 upregulation.

Importantly, this signature could potentially be applied to human samples to identify tumours with high/low BACH1 activity. However, it is still uncertain whether the genes identified using perturbation experiments such as BACH1 depletion or inhibition, serve as reliable surrogates for BACH1 basal levels or activity in human tissue, as this signature has not yet been tested in tissue. Further research is needed to determine the utility of this novel signature in identifying patients with high levels of BACH1, who could benefit most from BACH1 inhibitors.

## Supporting information

A549 wt vs BACH1 KO RNA seq data

A549 CHIP RNA-seq merged with Score

Homer list of known motifs A549 Bach1

## Funding

This work was supported by Cancer Research UK (C52419/A22869 to LV, DW and MH) and Ninewells Cancer Campaign (DK) and by the Swedish Research Council (2018-02318 and 2022-00971 to VIS, 2021-03138 to CW), the Swedish Cancer Society (23-3062 to VIS, 22-0612FE to CW), Assar Gabrielsson Research Foundation (to AAHP, DR, CW, and VIS), the Swedish Society for Medical Research (2018; S18-034 to VIS), the Knut and Alice Wallenberg Foundation, and the Wallenberg Centre for Molecular and Translational Medicine (to VIS).

## Authors Contribution

DK, DW, KXA, MH, AC, CM and DR conducted the cell biology experiments and KXA, AAHP and DR the bioinformatic analysis. DK, DW, KXA and CW were responsible for data analysis, figure preparation and statistical analysis. TW and EL provided resources and technical expertise. LV, VIS and CW had a leading contribution in the design of the study, and an active role in the discussion and interpretation of the whole dataset. LV wrote the original draft of the manuscript. All authors reviewed and edited the manuscript. Funding acquisition LV, VIS and CW. All authors take full responsibility for the work.

---

**Supplementary Figure S1.**
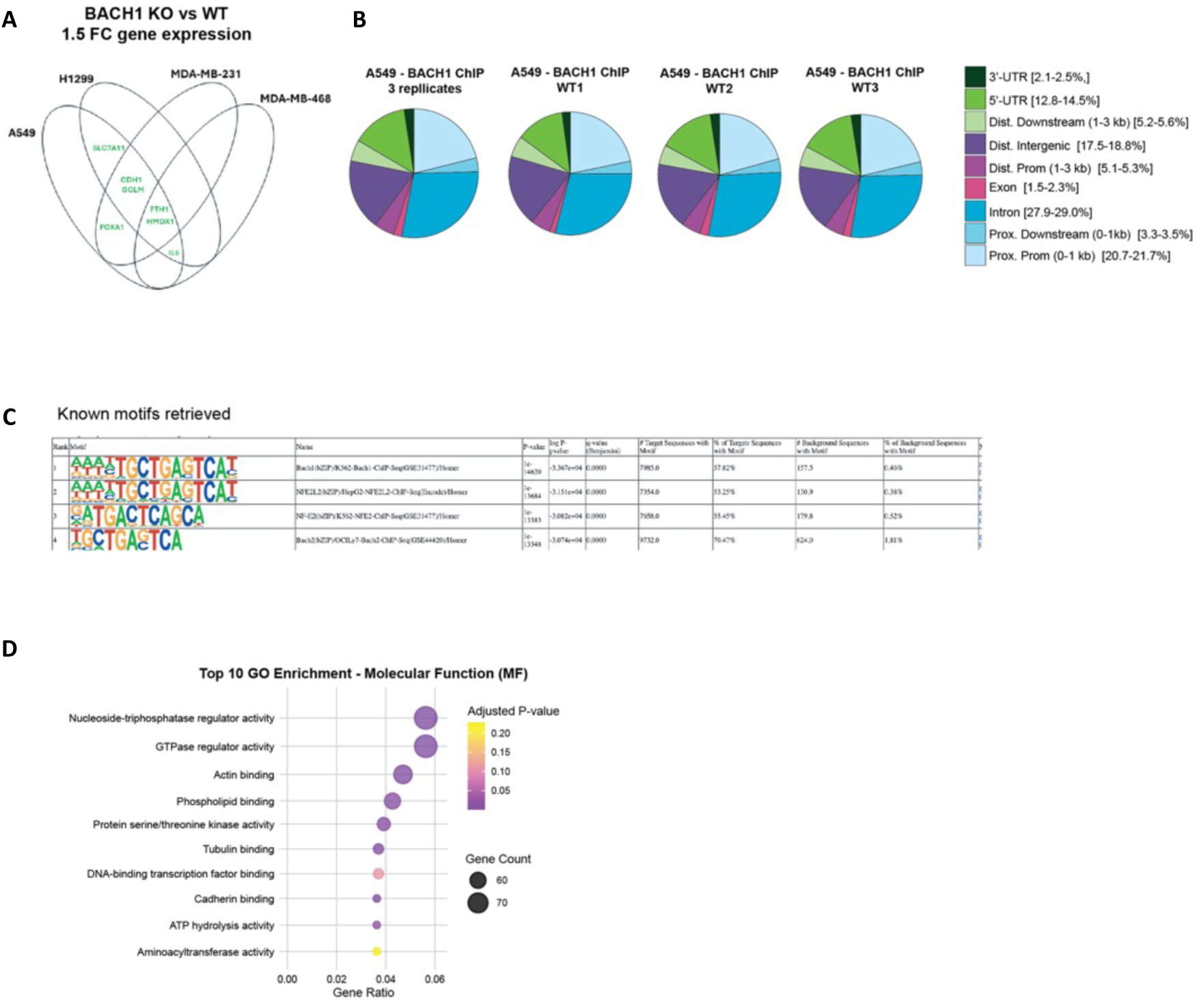
**A)** Venn diagram illustrating the overlap of differentially expressed BACH1 target genes upon BACH1 depletion using a 1.5-fold cutoff across A549, H1299, MDA-MB-231, and MDA-MB-468 cell lines. Genes such as *FTH1, GCLM, FOXA1, CDH1, SLC7A11*, and *IL6* showed partial overlap across some cell lines, while *HMOX1* remained the only gene shared among all four cell lines. **B)** Genomic distribution of BACH1 binding sites in A549 wild-type replicates (WT1, WT2, WT3) and the combined dataset (3 replicates). Binding sites show a preferential distribution in introns (28.5%), promoters within 1 kb of transcription start sites (21.7%), and intergenic regions (17.8%), demonstrating consistency across the replicates. **C)** Table of known motifs retrieved from HOMER analysis for BACH1 ChIP-Seq in A549 cells. Each motif is displayed with its sequence logo and associated gene. **D)** Top 10 Gene Ontology (GO) enrichment analysis for molecular functions (MF) performed on data integrating RNA-Seq and ChIP-Seq genes.

**Supplementary Figure S2.**
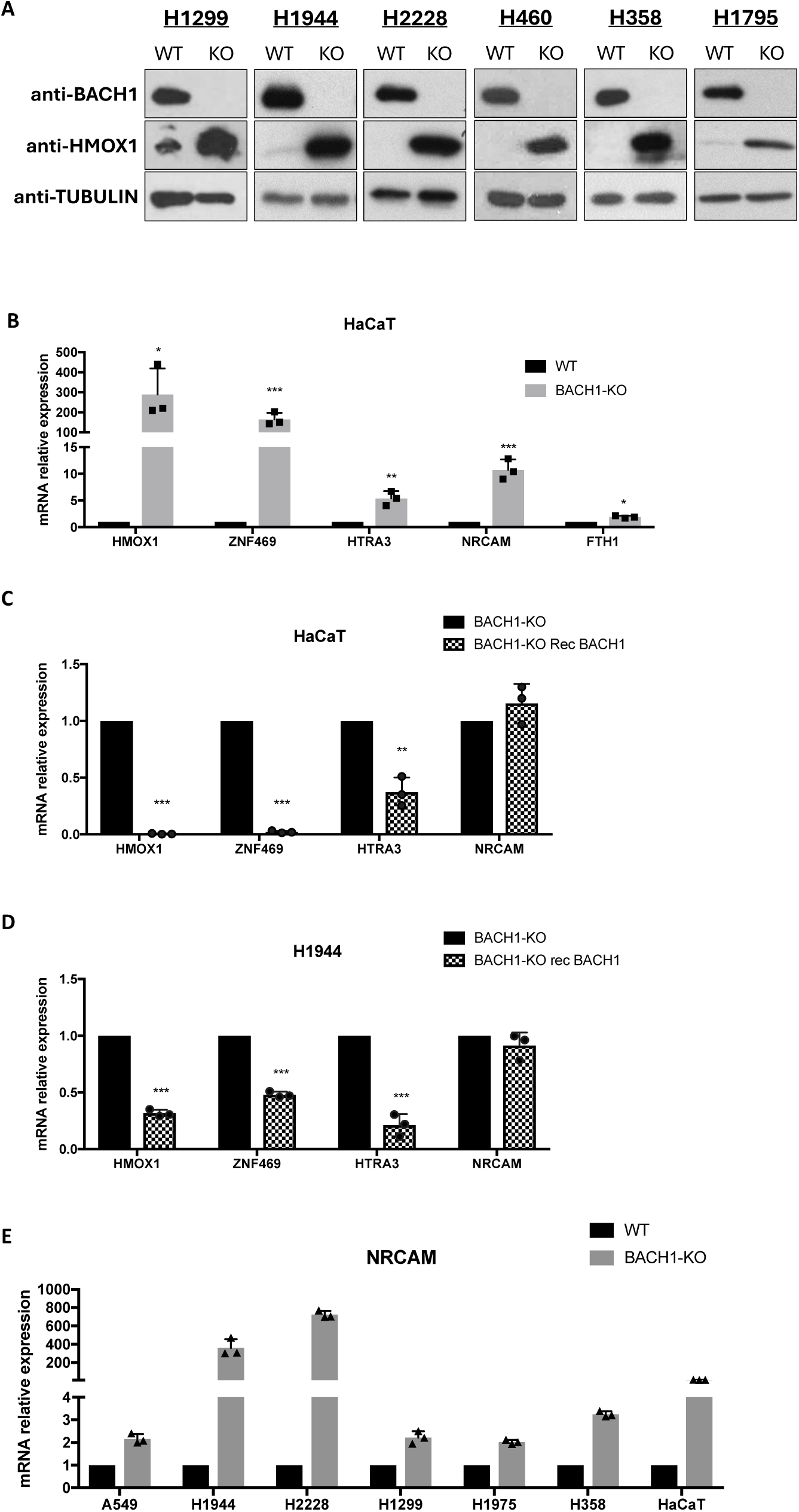
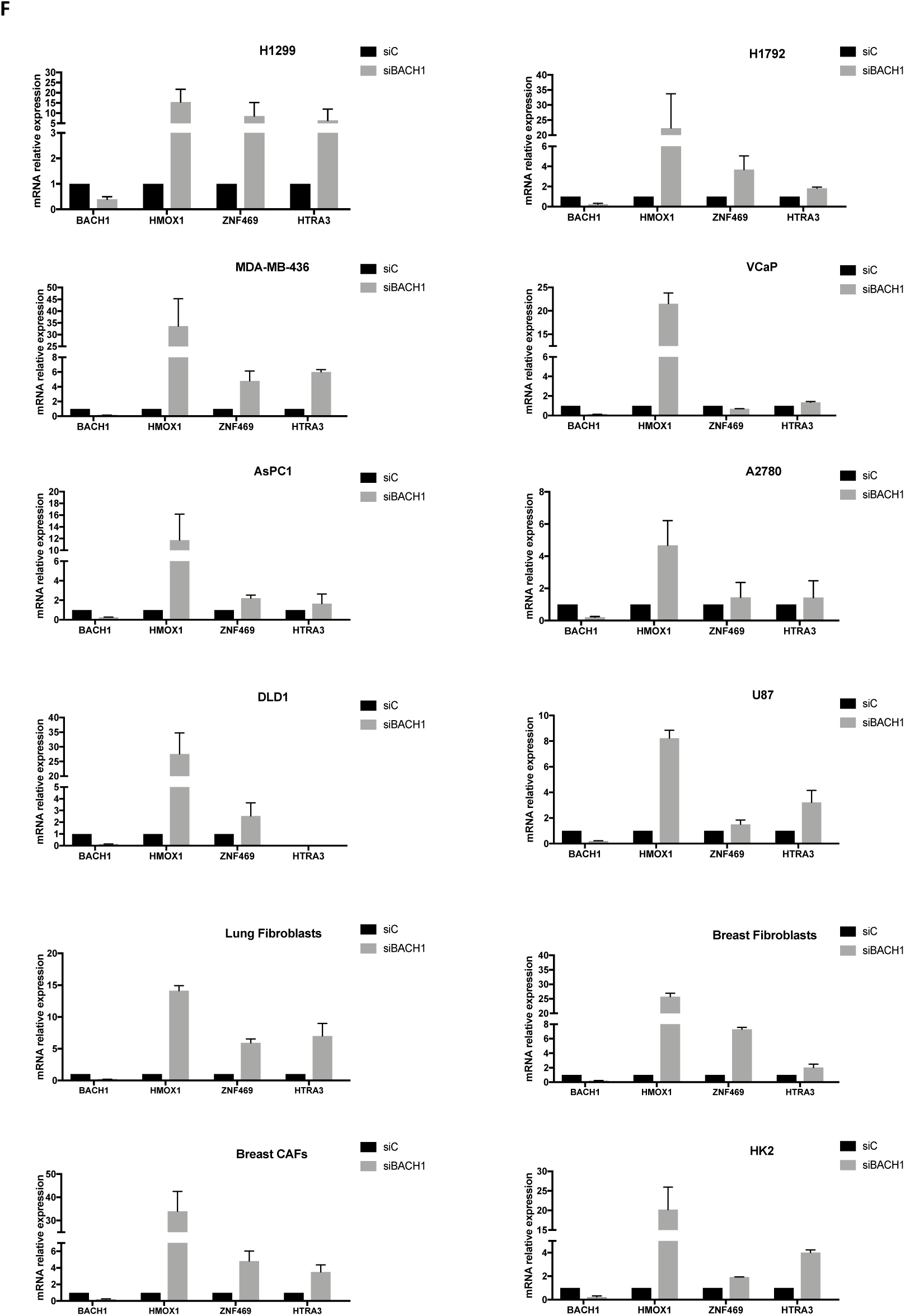
**A)** Lysates from a panel of control (WT) and BACH1-KO lung cancer cells were analysed by western blot for the indicated proteins. **B)** HaCaT control (WT) and BACH1-KO cells were lysed and mRNA levels of the indicated genes were analysed by real-time qPCR. Data (n=3) represent means± SD and are expressed relative to the WT sample. **C and D)** HaCaT BACH1-KO or H1944 BACH1-KO cells were infected with either GFP control or with BACH1-GFP virus (rec BACH1) and selected with puromycin (4 µgr/ml). 72 hours later cells were lysed, and mRNA levels of the indicated genes were analysed by real-time qPCR. Data (n=3) represent means± SD and are expressed relative to the BACH1-KO (GFP control) sample. **E)** A panel of control (WT) and BACH1-KO cell lines were compared for their expression of NRCAM. Data (n=3) represent means± SD and are expressed relative to the control sample. **F)** The indicated cell lines were transfected with either siControl (siC) or siBACH1. 48 hours later cells were lysed, and mRNA levels of the indicated genes were analysed by real-time qPCR. Data (n=2) represent means± SD and are expressed relative to the siControl sample.

**Supplementary Figure S3.**
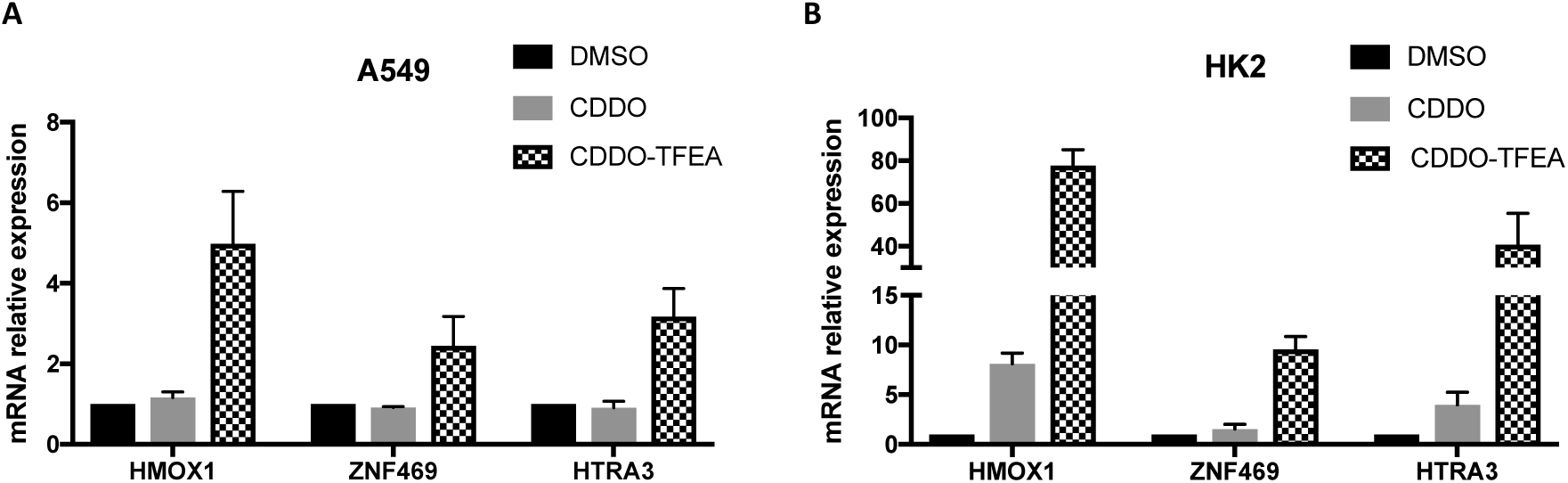
**A and B)** A549 cells (*left panel*) or HK2 cells (*right panel*) were treated with either DMSO, CDDO (100 nM) or CDDO-TFEA (100 nM) for 16 hours. After that, cells were lysed, and mRNA levels of the indicated genes were analysed by real-time qPCR. Data (n= 3) represent means± SD and are expressed relative to the DMSO treated sample.

**Supplementary Figure S4.**
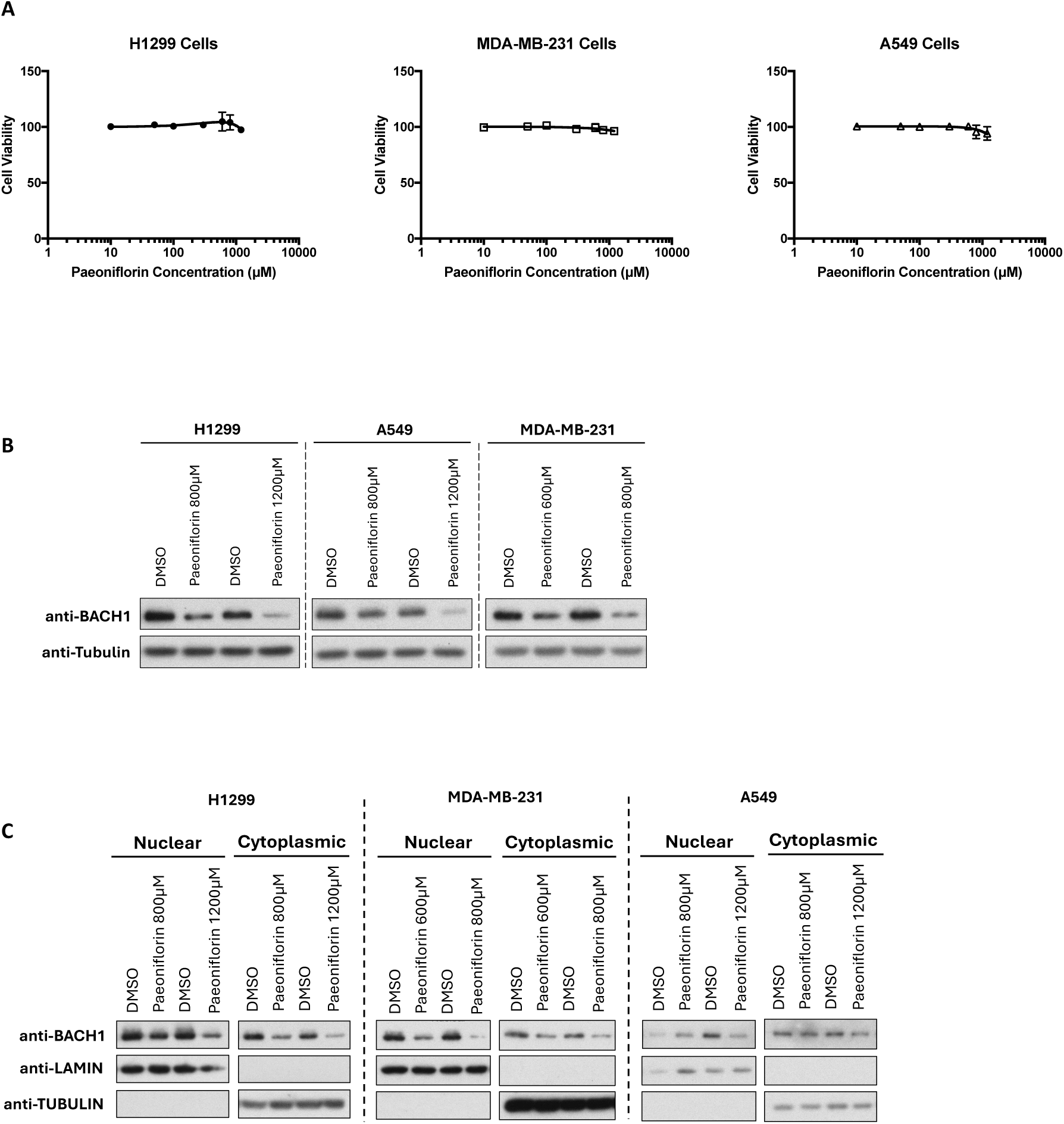
**A)** The indicated cell lines were treated with either DMSO or increasing concentrations of paeoniflorin as indicated. After 48 hours, viability was calculated relative to the DMSO-treated control using Alamar Blue. Data (n=3) represent means ± SD and are expressed relative to the DMSO treated sample. **B)** *Related to Fig. 4B*. Lung cancer cells (A549 and H1299) or breast cancer cells (MDA-MB-231) were treated with vehicle (DMSO) or with paeoniflorin at the indicated concentrations. After 16 hours cells were lysed, and samples were analysed by western blot with the indicated antibodies. The figure shows a representative blot. **C)** *Related to Fig. 4B*. Like in B) but cells were subjected to subcellular fractionation and separated in nuclear and cytoplasmic fractions. The figure shows a representative blot.

**Supplementary Figure S5.**
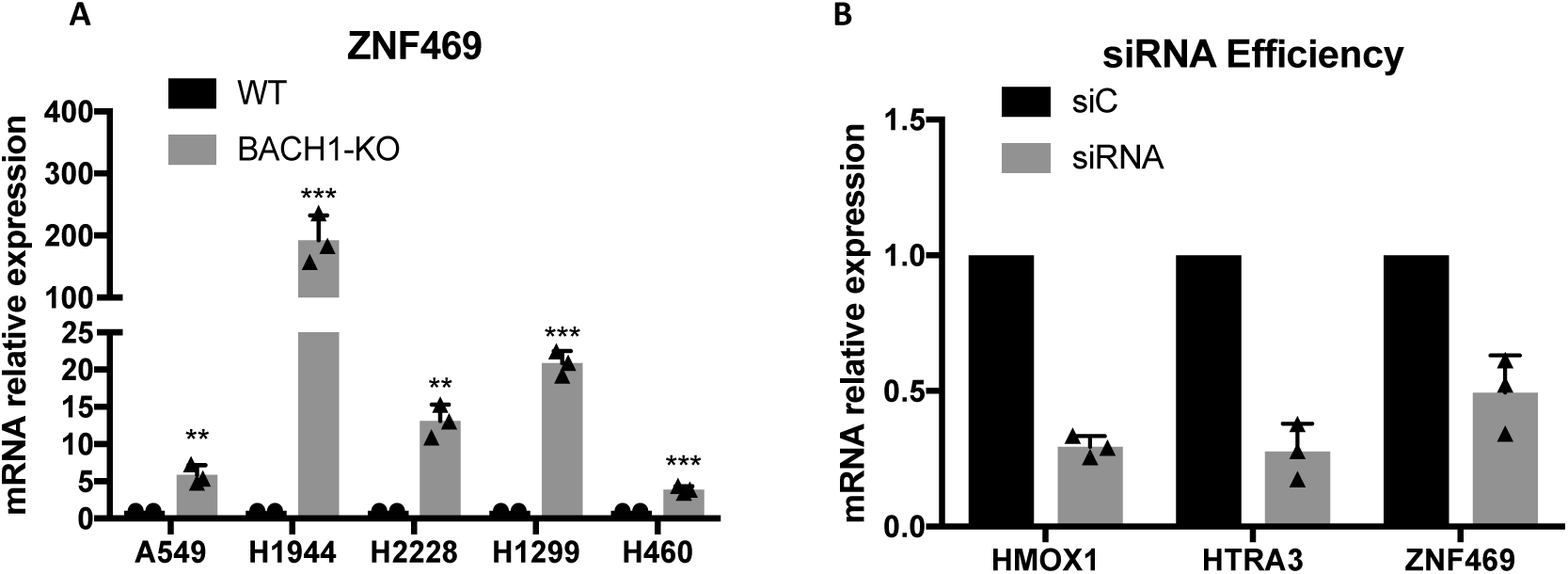
**A)** A panel of control (WT) and BACH1-KO lung cancer cell lines were compared for their expression of *ZNF469*. Data (n=3) represent means± SD and are expressed relative to the control sample. **B)** *Related to Figure 5C*. 10% of each sample was analysed by RT-PCR to confirm the efficiency of the silencing by comparing them against the siControl sample.

## Notes

### Competing Interest Statement

The authors have declared no competing interest.

