## Supplementary Material and Methods for "Identification of a BACH1 lung cancer signature: A novel tool for understanding BACH1 biology and identifying new inhibitors"

Taqman probes used:

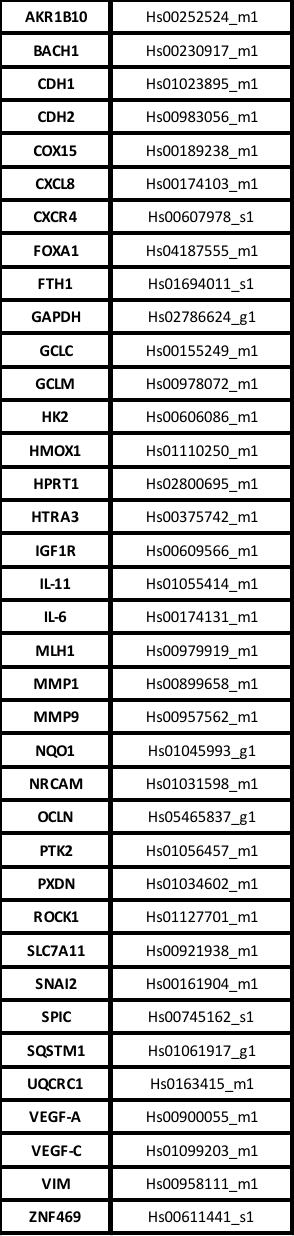

List of reported BACH1 targets tested:

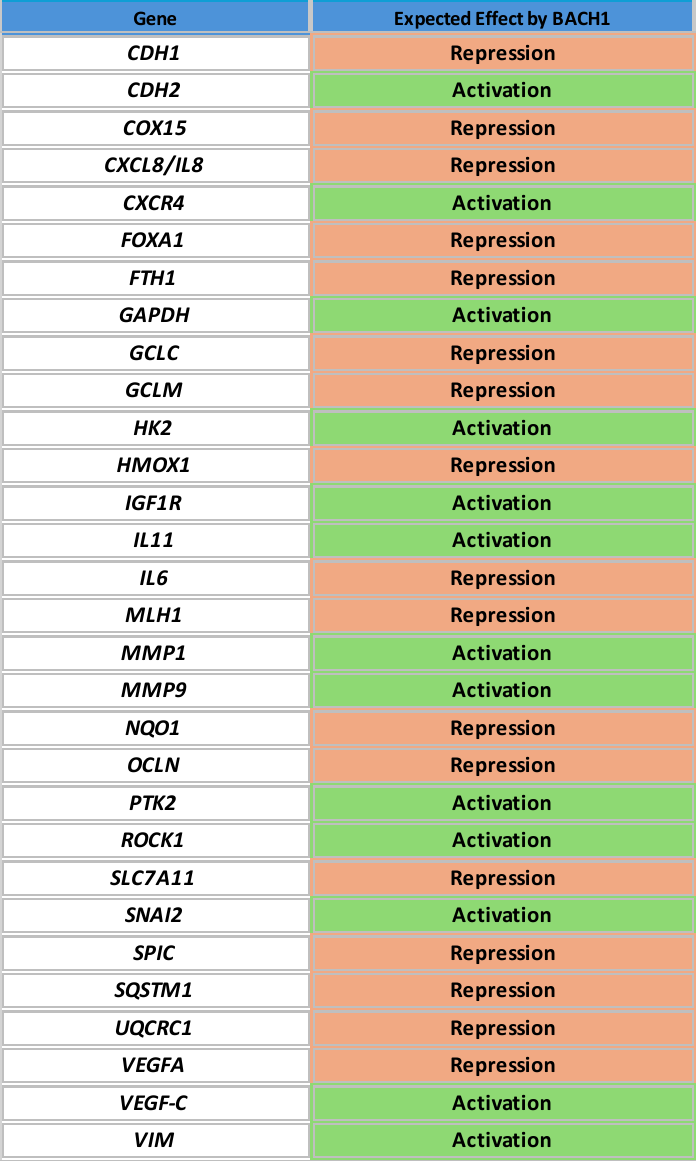
